# Fractional and Geometric Neural Dynamics: Investigating Intelligence-Related Differences in EEG Symmetry and Connectivity

**DOI:** 10.1101/2025.02.25.640025

**Authors:** Arturo Tozzi, Ksenija Jaušovec

**Author notes:** (corresponding author) 1155 Union Circle, #311427 Denton, TX 76203-5017 USA.

## Abstract

Understanding intelligence-related variations in electroencephalographic (EEG) activity requires advanced mathematical approaches capable of capturing geometric transformations and long-range dependencies in neural dynamics. These approaches may provide methodological advantages over conventional spectral and connectivity-based techniques by offering deeper insights into the structural and functional organization of neural networks. In this study, we integrate Clifford algebra, Noether’s theorem and fractional calculus to analyze EEG signals from high- and low-IQ individuals, looking for key intelligence-related differences in cortical organization. Clifford algebra enables the representation of EEG signals as multivectors, preserving both magnitude and directional relationships across cortical regions. Noether’s theorem provides a quantitative measure of symmetry properties linked to spectral features, identifying conserved functional patterns across distinct brain regions. Mittag-Leffler functions, derived from fractional calculus, characterize long-range dependencies in neural oscillations, allowing for the detection of memory effects and scale-invariant properties often overlooked by traditional methods. We found significant differences between high- and low-IQ individuals in geometric trajectories, hemispheric connectivity, spectral properties and fractional-order dynamics. High-IQ individuals exhibited increased spectral asymmetry, enhanced spectral differentiation, distinct geometric trajectories and greater fractional connectivity, particularly in frontal and central regions. In contrast, low-IQ individuals displayed more uniform hemispheric connectivity and heightened fractional activity in occipital areas. Mittag-Leffler fractional exponents further indicated that high-IQ individuals possessed more varied neural synchronization patterns. Overall, our multi-faceted approach suggests that intelligence-related neural dynamics are characterized by an asymmetric, functionally specialized and fractionally complex cortical organization. This results in significant differences in network topology, efficiency, modularity and long-range dependencies.

## INTRODUCTION

Intelligence has been associated with distinct electroencephalographic (EEG) neural patterns, yet the underlying mechanisms remain a subject of investigation (Thatcher et al., 2005; Friedman et al., 2019). The study of intelligence-related differences in EEG signals has traditionally relied on spectral, time-frequency and connectivity analyses (Chen et al., 2023; Ignatious et al., 2023). Conventional approaches such as Fourier and wavelet transforms, coherence measures and graph-theoretic network analyses have provided critical insights into brain function but lack the mathematical depth to fully capture neural relationships (Sitnikova et al., 2009; San-Segundo et al., 2019). We argue that mathematical frameworks integrating algebraic, geometric, and fractional dynamics provide an alternative perspective by capturing spatial dependencies and characterizing long-range memory effects in neural activity. Among these approaches, Clifford algebra has been used to encode EEG signals as multivectors, preserving both magnitude and directional relationships across cortical regions—an advantage not achievable with conventional spectral methods (Zhang et al., 2023). Clifford algebra-based EEG transformations allow for the preservation of geometric properties enabling novel trajectory-based comparisons. Noether’s theorem, which relates system symmetries to conserved quantities, has been used to assess functional and spectral balance in cortical microcircuitry, revealing conserved functional patterns across different cortical regions (Bilteanu et al., 2017). Meanwhile, fractional calculus — particularly Mittag-Leffler function analysis— may extend EEG signal characterization beyond integer-order models, allowing for the detection of memory effects and scale-invariant properties often overlooked by standard methods (Atanackovic et al., 2011; Turalska and West, 2018). Functional order analysis provides an additional layer of insight by quantifying fractional dynamics and long-range dependencies (García-Raffi and Torrano, 2021).

Together, these multi-faceted approaches may enable a refined investigation into intelligence-related brain dynamics extending beyond pairwise spectral differences, allowing for a richer assessment of interregional dependencies. We conjecture that our approach might reveal systematic differences in EEG structure, with high-IQ individuals exhibiting greater frontal asymmetry, enhanced connectivity in integrative brain areas and increased fractional-order complexity in neural synchronization. Low-IQ individuals, in contrast, might display more uniform hemispheric connectivity, potentially indicative of different cognitive resource allocation strategies.

In sum, given the limitations of traditional EEG methods, a hybrid approach that considers Clifford algebra, Noether’s theorem and fractional network measures holds promise for more comprehensive analyses of intelligence. We will proceed as follows. The next section outlines the methodology, detailing data acquisition, preprocessing and the mathematical frameworks applied to EEG signals. We then present our results, followed by an interpretation of the findings in the context of intelligence-related neural organization. Finally, we conclude with a discussion on the broader implications and future directions of this research.

## MATERIALS AND METHODS

Our retrospective study builds on prior research by Jaušovec and Jaušovec (2001; 2003; 2005; 2010) and later advancements with Tozzi et al. (2021a; 2021b), honoring Norbert Jaušovec’s contributions. EEG data were collected from 10 right-handed participants (mean age: 19.8 years; SD = 0.9; males: 4), divided into High-IQ (IQ SD = 127) and Low-IQ (IQ SD = 87) groups based on standardized test scores during auditory and visual oddball tasks. EEG signals were recorded using a 64-channel system with electrodes placed according to the 10–20 system. Data epochs were segmented into non-overlapping 2-second windows and trials contaminated with excessive noise were excluded based on an amplitude threshold exceeding ±100 µV. Standardization was performed by normalizing each EEG signal relative to its mean and standard deviation across trials, ensuring comparability between participants.

### Clifford algebra

Clifford algebra was used to transform EEG signals into multivector representations that preserve spatial and directional relationships between cortical regions (Acus and Dargys, 2024). Each EEG signal *x*_*i*_ (*t*) from electrode *i* was mapped onto a Clifford multivector Ψ(*t*), expressed as

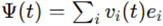

where *v*_*i*_ (*t*) is the voltage amplitude and *e*_*i*_ represents a basis vector of a Clifford algebra. The choice of Clifford space dimensionality depended on the number of selected EEG channels. In our study, a three-dimensional Clifford space *Cl*(3, 0) was chosen to encode signals from three representative electrode locations (FP1, P3, O1). These three electrodes were selected due to their known involvement in cognitive functions and information processing. The transformed EEG signals were then analyzed geometrically by computing trajectory deviations across trials. The Clifford Fourier Transform (CFT) was used to extract spectral features within the multivector space (Monaim and Fahlaoui, 2024). Given a time-dependent multivector Ψ(*t*), its frequency-domain representation was obtained as

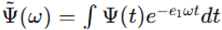

where *e*_1_ is an imaginary unit in Clifford space. Statistical comparisons of Clifford trajectories were performed using paired t-tests to determine significant differences in neural geometry between high and Low-IQ groups.

### Noether’s theorem

Noether’s theorem was used to quantify symmetry in functional connectivity. Functional connectivity was evaluated using Pearson correlation coefficients between electrode pairs, computed as

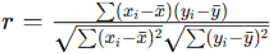

where *x*_*i*_ and *y*_*i*_ represent EEG signals from different electrodes and 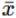 and 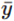 are their means over time. Connectivity matrices were constructed for each participant, with a focus on left-right hemisphere electrode pairs (FP1–FP2, F3–F4, C3–C4, P3–P4, O1–O2) to assess hemispheric symmetry. Symmetry indices were calculated as

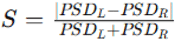

where *PSD*_*L*_ and *PSD*_*R*_ are power values from corresponding left and right hemisphere electrodes. Spectral asymmetry was assessed by performing a Fourier transform on each EEG channel and comparing power distributions between hemispheres (Atiyah et al., 1975). Welch’s t-tests were applied to compare connectivity and spectral symmetry indices between high and Low-IQ groups.

To assess the relationship between frequency band power and hemispheric symmetry, correlation analyses were performed for each frequency band. Power spectral densities (PSDs) were computed to assess frequency distributions in delta, theta, beta and alpha bands (Dressler et al., 2004; Redwan et al., 2024). Pearson correlation coefficients were then calculated between frequency band power and hemispheric symmetry indices to determine whether specific oscillations were associated with neural organization patterns.

To evaluate the temporal stability of EEG signals, stationarity tests were conducted using the Augmented Dickey-Fuller (ADF) test, which determines whether a time series exhibits long-term trends or remains stable over time (Dao and Staszewski, 2021). ADF p-values below 0.05 indicated significant stationarity, suggesting a consistent neural state.

### Fractional derivative values

Fractional-order derivatives were computed to characterize the temporal dynamics of EEG signals, as conventional differentiation does not account for memory effects or long-range dependencies inherent in neural activity. The Caputo fractional derivative was chosen, defined as

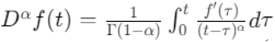

where *α* represents the fractional order and Γ denotes the Gamma function. The parameter *α* was optimized based on the signal’s spectral properties to capture its underlying fractional nature effectively. Numerical implementation was carried out using the Grünwald-Letnikov approximation, which approximates the integral definition with a discrete summation:

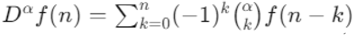

where 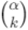 represents the binomial coefficient extended to non-integer orders. This method provided a means of assessing long-range dependencies in EEG data, differentiating between stationary and non-stationary signal properties. The fractional derivative values were extracted for each EEG channel and used in subsequent statistical analyses to identify group-wise differences in neural activity. These calculations formed the basis for evaluating the Mittag-Leffler function’s role in neural dynamics, establishing a mathematical framework for comparing cognitive groups.

### Fractional connectivity and fractional clustering coefficients

To examine functional connectivity, correlation-based adjacency matrices were constructed by computing pairwise cross-correlations between EEG channels. Cross-correlation was defined as

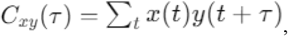

where *x*(*t*) and *y*(*t*) are EEG signals from two channels and *τ* represents the lag. A sliding window approach was used to ensure robust estimation of functional connectivity over time, with overlapping one-second segments applied to account for dynamic variations in brain activity.

To integrate fractional-order concepts into network analysis, fractional connectivity and fractional clustering coefficients were computed. Fractional connectivity was derived by modifying edge weights using a power-law transformation, where each connection strength *W*_*ij*_ was raised to a fractional exponent *γ*, yielding 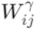: The mean fractional connectivity was then calculated by averaging these transformed values, providing a metric sensitive to longrange dependencies in neural networks.

The fractional clustering coefficient was computed using a modified version of the standard clustering measure, incorporating the fractional-weighted edges to assess local connectivity structures. This was defined as

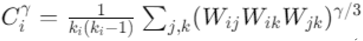

where *k*_*i*_ is the degree of node *i*. These fractional measures allowed for a more nuanced characterization of network topology, enabling differentiation between classical small-world organization and fractional network configurations.

The ability to quantify connectivity and clustering using fractional-order metrics provided insights into how neural interactions vary across cognitive groups, extending conventional graph-theoretic approaches.

### Mittag-Leffler fractional exponents

Mittag-Leffler fractional exponents were estimated to describe the scaling behavior of connectivity weights within the network (García-Raffi and Torrano, 2021). The Mittag-Leffler function, defined as

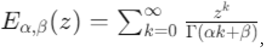

generalizes exponential decay and captures memory effects in complex systems (Tarasov 2018). To estimate the effective fractional exponent, the probability distribution of connectivity weights was examined in log-log space and a power-law function

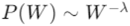

was fitted to the data. The exponent λ was extracted using least-squares fitting, providing a measure of the heterogeneity in connectivity distributions. Higher values of λ indicated a broader range of connectivity strengths, reflecting a more diverse interaction pattern across EEG channels. This Mittag-Leffler-based analysis was used to assess differences in the scaling properties of functional networks between groups, serving as a crucial parameter for understanding cognitive variability.

Statistical analyses were performed to compare fractional network measures between high and Low-IQ groups. Oneway analysis of variance (ANOVA) was conducted to test for significant differences in fractional connectivity, fractional clustering and Mittag-Leffler exponents across groups. The Bonferroni correction was applied to account for multiple comparisons and control the false discovery rate.

### Tools

Computational analyses were implemented using Python, utilizing NumPy for numerical computations, SciPy for statistical analysis, Matplotlib for visualization and the MNE-Python toolbox for EEG preprocessing. Clifford algebra computations were performed using the ClPy library and Fourier transforms were applied using SciPy’s signal processing module.

## RESULTS

Significant differences in geometric trajectories, hemispheric connectivity, spectral properties and fractional-order dynamics were found between high and Low-IQ individuals, with High-IQ individuals exhibiting increased Clifford algebra-based frontal variability, greater frontal asymmetry and enhanced spectral differentiation.

### Clifford algebra

Clifford algebra-based transformations displayed distinct EEG trajectory patterns, with High-IQ individuals exhibiting increased geometric variability in the frontal region, while Low-IQ individuals exhibited more constrained neural trajectories (**Figure 1A**). Statistical comparisons of Clifford components indicated significant differences in frontal representations between groups (p < 0.001), whereas no significant differences were found in parietal and occipital components (**Figure 1B**). Temporal stationarity analyses using the Augmented Dickey-Fuller test indicated that EEG signals in High-IQ individuals were more stationary (p < 0.05), suggesting greater consistency in neural oscillations.

**Figure 1.**
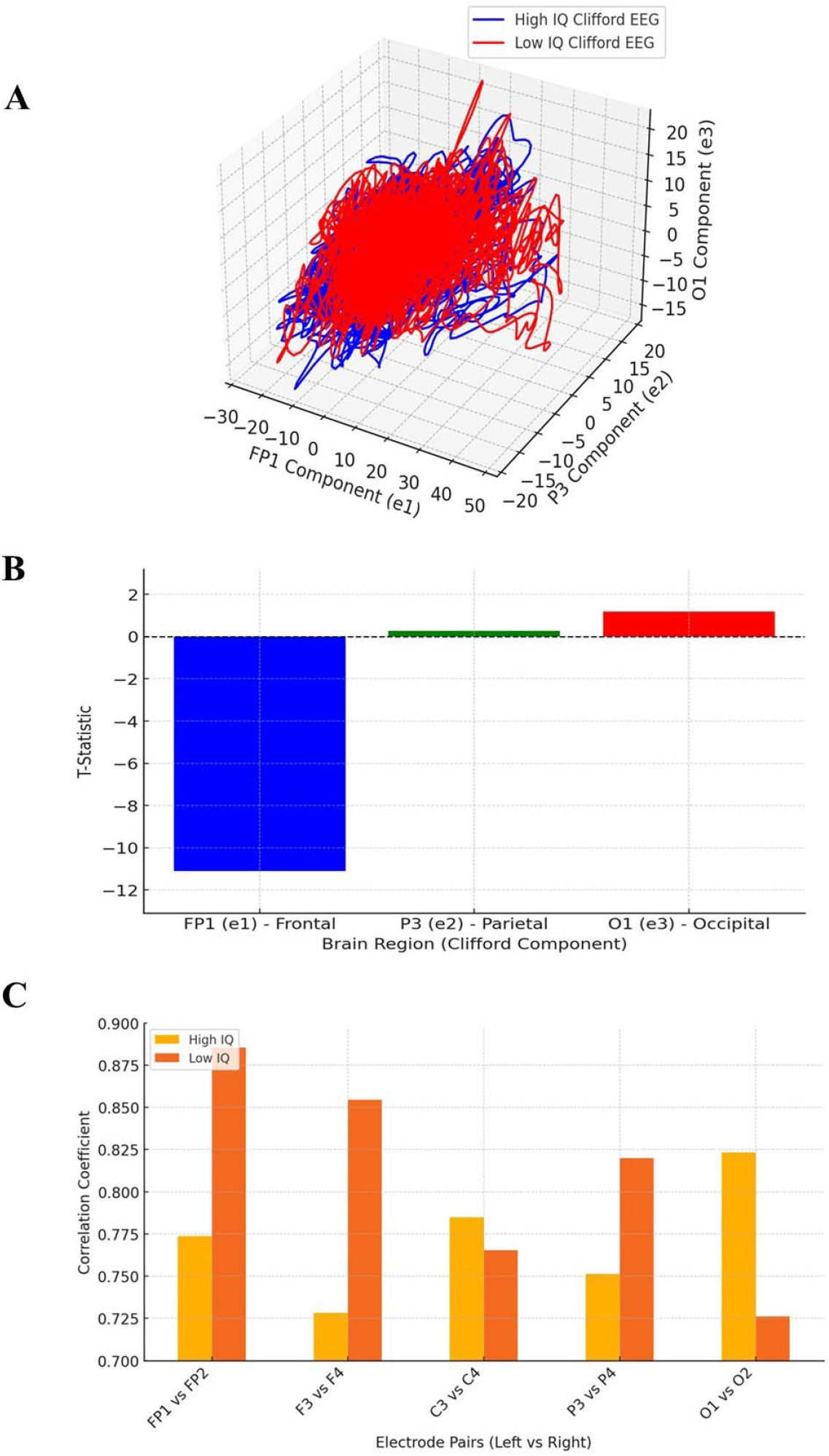
**A.**Clifford EEG Signal representation for High vs. Low-IQ groups. The blue and red lines represent average EEG trajectories in Clifford space for high and Low-IQ individuals, respectively. Differences in shape, spread and oscillatory patterns provide insight into IQ-related brain activity variations. **Figure 1B**. Statistical differences in Clifford EEG components. Significant differences were observed in the frontal region, while parietal and occipital components showed no substantial variation. The dashed line at zero indicates the statistical significance threshold. **Figure 1C**. Hemispheric symmetry differences across brain regions. Low-IQ individuals (green bars) exhibit greater symmetry in prefrontal, frontal and parietal regions, whereas High-IQ individuals (blue bars) exhibit increased asymmetry, particularly in FP1-FP2 and F3-F4. The occipital region (O1-O2) is more symmetric in High-IQ individuals, suggesting enhanced visual-spatial processing.

Spectral power analysis using the Clifford Fourier Transform showed that High-IQ individuals exhibited significantly higher alpha (p < 0.001) and beta (p = 0.0067) power in the frontal region compared to Low-IQ individuals, suggesting different neural activation patterns. Power spectral densities showed that High-IQ individuals had significantly higher theta (p = 0.0065) and beta (p = 0.0067) power, while Low-IQ individuals exhibited relatively stronger delta oscillations.

### Noether’s theorem

Connectivity analysis based on Noetherian symmetry principles and further analyses of spectral symmetry revealed significant differences in hemispheric organization. High-IQ individuals displayed greater asymmetry in frontal connectivity (FP1-FP2, p < 0.001) and enhanced symmetry in occipital connectivity (O1-O2, p < 0.001), while Low-IQ individuals maintained more uniform hemispheric connectivity, particularly in prefrontal and parietal regions (**Figure 1C**).

### Fractional dynamics

Statistical analysis of Mittag-Leffler fractional derivatives revealed distinct patterns of neural activity between high- and low-IQ individuals. High-IQ individuals consistently exhibited higher fractional values in frontal (FP1, F4) and central (C3) regions, while low-IQ individuals showed increased fractional activity in the occipital (O1) area (**Figure 2A**). Notably, the frontal regions displayed significantly lower fractional-order dynamics in the low-IQ group, suggesting that high-IQ individuals engage in neural processes with stronger memory effects and long-range dependencies, potentially supporting cognitive flexibility and problem-solving.

**Figure 2.**
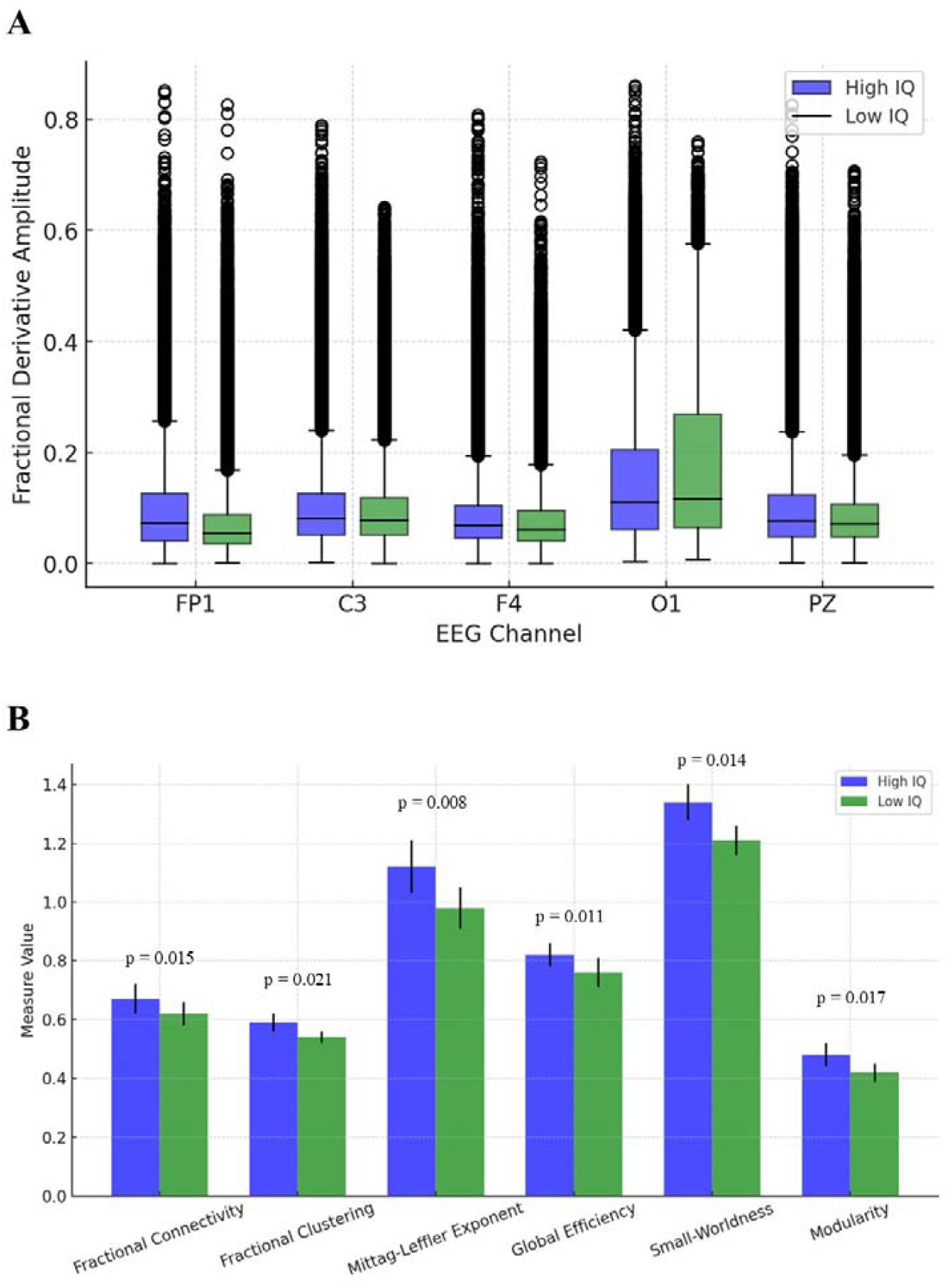
**A**. Boxplot comparison illustrating key differences in fractional dynamics between High-IQ (blue) and Low-IQ (green) groups. The variations in distributions suggest significant differences in Mittag-Leffler function characteristics. Statistical analysis confirms that these distinctions are significant, with **p < 0.0001** across all analyzed channels. **Figure 2B**. Comparison of fractional network measures between high and Low-IQ groups. Bars represent mean values for each group, with error bars indicating standard deviations. High-IQ individuals exhibit significantly greater fractional connectivity, clustering, Mittag-Leffler exponent, global efficiency, small-worldness and modularity compared to Low-IQ individuals. One-way ANOVA confirmed systematic distinctions in neural structure and connectivity patterns across groups.

Beyond localized effects, significant differences were also observed in central and parietal channels, indicating that intelligence-related variability extends across broader network properties rather than being confined to specific brain regions. The occipital region exhibited an inverse trend, with low-IQ individuals displaying higher Mittag-Leffler fractional activity, possibly reflecting distinct neural connectivity patterns or alternative sensory integration strategies. Furthermore, greater variability in fractional activity was observed in some channels within the low-IQ group, as indicated by wider distribution ranges. This suggests increased dispersion in fractional-order neural activity, which may correspond to less stable or less efficient network dynamics.

### Fractional network measures

The comparison of fractional network measures between high and Low-IQ groups is illustrated in **Figure 2B**. Fractional connectivity exhibited significant differences between groups, with High-IQ individuals showing an higher mean fractional connectivity. Similarly, fractional clustering coefficients were higher in the High-IQ group. Mittag-Leffler fractional exponents demonstrated a broader distribution in the High-IQ group, with a mean exponent value lower in the Low-IQ group. Further, High-IQ individuals exhibited greater variability in fractional connectivity values. Weighted global efficiency was significantly higher in the High-IQ group, suggesting differences in the efficiency of information transfer across neural networks. This set of findings highlights a more complex and distributed network topology in individuals with High-IQ, underlining the relevance of fractional measures in characterizing cognitive differences. Small-worldness, defined as the ratio of clustering coefficient to characteristic path length, was significantly greater in the High-IQ group. This suggests that High-IQ individuals exhibit a more optimized network structure, balancing local clustering with global efficiency. Additionally, edge weight distribution followed a broader power-law decay in the High-IQ group, consistent with more heterogeneous connectivity patterns.

The Mittag-Leffler-based scaling exponent analysis further corroborated these findings, as High-IQ individuals demonstrated greater deviation from exponential connectivity decay, indicating a higher degree of long-range dependencies in neural interactions. These variations were particularly evident in the frontal and occipital regions, where High-IQ individuals displayed enhanced connectivity strengths. Differences in network modularity also emerged, with High-IQ individuals exhibiting higher modularity scores, suggesting a more functionally segregated yet highly interactive network.

Overall, these findings suggest that fractional network properties, including connectivity, clustering and Mittag-Leffler exponents, may distinguish high and Low-IQ groups in a statistically significant manner.

Summarizing, our findings, confirmed through Clifford algebra transformations, spectral analysis and Noetherian symmetry principles, suggest that High-IQ individuals exhibit greater frontal EEG asymmetry, increased spectral differentiation and distinct connectivity patterns, greater geometric variability, greater frontal asymmetry, enhanced connectivity in integrative regions and increased long-range dependencies, while Low-IQ individuals displayed more uniform hemispheric connectivity and heightened activity in occipital areas. These findings suggest that intelligence-related neural dynamics are characterized by asymmetric, functionally specialized and fractionally complex cortical organization.

## CONCLUSIONS

We investigated intelligence-related differences in EEG signals by combining multiple mathematical approaches such as Clifford algebra, Noether’s theorem and fractional network analysis. Functional connectivity analysis revealed that high-IQ individuals exhibited a more differentiated neural network organization that was characterized by greater variability in interregional interactions and enhanced symmetrical connectivity in occipital regions. Fractional-order analyses demonstrated that High-IQ individuals exhibited greater long-range dependencies, with significantly higher Mittag-Leffler fractional exponents and clustering coefficients, supporting the presence of more complex and efficient neural architecture. This suggests that cognitive ability may be linked to distinct neural processing strategies, with High-IQ individuals displaying more complex fractional-order behavior in regions associated with executive function and working memory. In turn, Low-IQ individuals exhibited heightened activity in the occipital cortex, which may correspond to alternative visual processing mechanisms.

High-IQ individuals displayed stronger Mittag-Leffler dynamics in regions linked to complex cognitive processing, while Low-IQ individuals displayed greater activity in visual and sensory processing areas. These differences align with theories of cognitive efficiency and regional specialization, where High-IQ brains may rely more on integrative network structures, while Low-IQ brains may exhibit increased reliance on localized sensory processing. Fractional-order dynamics showed that the frontal and central brain regions displayed more Mittag-Leffler behavior in High-IQ individuals. The occipital region presented an inverse trend, where Low-IQ individuals showed greater fractional activity, suggesting distinct processing dynamics. Taken together, these findings suggest that intelligence-related brain function may be reflected in both geometric transformations and fractional connectivity properties.

The integration of Clifford algebra, Noether’s theorem and Mittag-Leffler function-based fractional calculus stands for a novel approach to EEG analysis which offers methodological advantages over conventional spectral and connectivity-based techniques. Clifford algebra preserves magnitude and directional relationships between EEG signals, allowing for a richer representation of neural trajectories’ geometric transformations. Noether’s theorem introduces a principled framework for assessing conserved properties in brain networks, enabling the quantification of functional symmetry and intelligence-related differences in interregional connectivity patterns. Fractional network measures extend beyond traditional graph-theoretic approaches by capturing long-range dependencies and scale-invariant properties in neural oscillations. This may reveal subtle variations in brain dynamics that conventional EEG methodologies fail to detect and that are often overlooked in standard time-frequency analyses.

Compared to traditional EEG analysis techniques, our approach offers several advantages. Spectral analysis methods such as Fourier and wavelet transforms are widely used to decompose EEG signals into frequency components but do not retain spatial or directional relationships between electrodes (Yuan et al., 2018; Daud and Sudirman, 2022). Functional connectivity metrics, including coherence and phase synchronization, assess interregional interactions but often rely on pairwise comparisons that may overlook global network dependencies (Miskovic and Keil, 2015; Abdullateef et al., 2022). Graph-theoretic measures capture aspects of brain topology but do not explicitly incorporate long-range dependencies or fractional-order dynamics. In contrast, our combination of Clifford algebra, Noether’s theorem and fractional network analysis preserves both local signal variations and global connectivity patterns. The inclusion of fractional calculus further enhances sensitivity to memory effects and non-stationary dynamics (Chen and Wang, 2020).

Our methodological framework has potential applications in different fields. EEG-based cognitive profiling could benefit from Clifford algebra transformations to characterize individual differences in neural geometry. The integration of Noether’s theorem into EEG analysis could provide a means of quantifying hemispheric symmetry, which has implications for studying neurodevelopmental disorders. Fractional network measures could be employed in neurodegenerative research to monitor cognitive decline, as long-range dependencies and connectivity scaling properties may serve as biomarkers for early-stage disorders. The integration of Noether’s theorem into EEG analysis may also contribute to the study of neurodevelopmental and neuropsychiatric disorders, where alterations in hemispheric symmetry have been linked to conditions such as autism spectrum disorder, schizophrenia and dyslexia (Saugstad, 1999; Gage et al., 2009; Guo et al., 2013; Perkins et al., 2014; He et al., 2023). Still, brain-computer interface technologies may enhance classification accuracy by utilizing fractional-order EEG features, which incorporate long-range memory effects into machine learning models (Tarasov 2018).

Beyond these applications, our approach suggests testable experimental hypotheses, such as: a) whether interventions like transcranial stimulation can modulate EEG symmetry in ways consistent with Noetherian principles; b) whether cognitive training enhances fractional connectivity measures over time; c) whether neural efficiency, as quantified by Clifford algebra-based geometric transformations, correlates with cognitive performance in different task-based EEG paradigms; d) whether the observed intelligence-related differences in EEG connectivity persist over time, particularly in relation to neuroplasticity and cognitive aging. These experimental extensions reinforce the potential of our framework in broadening the scope of nervous oscillations’ research.

Despite its contributions, our study has several limitations. The sample size may limit the generalizability of findings and the robustness of observed differences. The computational complexity of Clifford algebra transformations presents a potential challenge, as real-time applications may require optimization to improve processing efficiency. While Noether’s theorem provides a theoretical basis for assessing neural symmetry, its direct application to EEG data for the study of intelligence requires further validation. Still, the selection of fractional exponent parameters was optimized based on empirical signal properties, but individual variations in neural dynamics may necessitate adaptive algorithms to improve robustness. Multimodal neuroimaging approaches, such as functional magnetic resonance imaging, could be incorporated to enhance spatial resolution and validate EEG-based symmetry findings.

In conclusion, we suggest that intelligence-related differences in EEG signals may be characterized using a combination of Clifford algebra, Noether’s theorem and fractional network analysis. By integrating these advanced mathematical frameworks, a novel perspective on cognitive variability could be provided, highlighting the role of symmetry, geometric transformations and fractional-order properties in brain function.

## DECLARATIONS

### Ethics approval and consent to participate

This research does not contain any studies with human participants or animals performed by the Author.

### Consent for publication

The Author transfers all copyright ownership, in the event the work is published. The undersigned author warrants that the article is original, does not infringe on any copyright or other proprietary right of any third part, is not under consideration by another journal and has not been previously published.

## Availability of data and materials

All data and materials generated or analyzed during this study are included in the manuscript. The Author had full access to all the data in the study and took responsibility for the integrity of the data and the accuracy of the data analysis.

## Acknowledgements

none.

## Authors’ contributions

The Author performed: study concept and design, acquisition of data, analysis and interpretation of data, drafting of the manuscript, critical revision of the manuscript for important intellectual content, statistical analysis, obtained funding, administrative, technical and material support, study supervision.

## Declaration of generative AI and AI-assisted technologies in the writing process

During the preparation of this work, the author used ChatGPT 4o to assist with data analysis and manuscript drafting and to improve spelling, grammar and general editing. After using this tool, the author reviewed and edited the content as needed, taking full responsibility for the content of the publication.

## REFERENCES

1) Abdullateef, S., B. Jordan, V. Rae, A. McLellan, J. Escudero, V. Nenadovic and T. Lo. “Quantitative Detection of Seizures with Minimal-Density EEG Montage Using Phase Synchrony and Cross-Channel Coherence Amplitude in Critical Care.” Annual International Conference of the IEEE Engineering in Medicine and Biology Society 2022 (July 2022): 259–262. 10.1109/EMBC48229.2022.9871595.

2) Acus, A. and A. Dargys. “Multivector (MV) Functions in Clifford Algebras of Arbitrary Dimension: Defective MV Case.” arXiv preprint 2412.05730 [math-ph], 2024. 10.48550/arXiv.2412.05730.

3) Atanackovic, Teodor M., Stevan Pilipovic, Bogoljub Stankovic and Dušan Zorica. 2011. “Fractional Calculus with Applications in Mechanics: Wave Propagation, Impact and Variational Principles.” Mathematical Problems in Engineering 2011: 298628. 10.1155/2011/298628.

4) Atiyah, M. F., V. K. Patodi and I. M. Singer. “Spectral Asymmetry and Riemannian Geometry. I.” Mathematical Proceedings of the Cambridge Philosophical Society 77, no. 1 (1975): 43–69.

5) Bilteanu, L., M. F. Casanova and I. Opris. “Symmetry and Noether Theorem for Brain Microcircuits.” In The Physics of the Mind and Brain Disorders, edited by I. Opris and M. F. Casanova, vol. 11, Springer Series in Cognitive and Neural Systems. Cham: Springer, 2017. 10.1007/978-3-319-29674-6_6.

6) Chen, Y. M. and J. R. Wang. 2020. “A High-Order Compact Finite Difference Scheme for the Time Fractional Black–Scholes Model.” Journal of Computational Physics 409: 109333. 10.1016/j.jcp.2020.109333.

7) Chen, Di, Haiyun Huang, Xiaoyu Bao, Jiahui Pan and Yuanqing Li. “An EEG-Based Attention Recognition Method: Fusion of Time Domain, Frequency Domain and Non-Linear Dynamics Features.” Frontiers in Neuroscience 17 (July 12, 2023). 10.3389/fnins.2023.1194554.

8) Cui, G., X. Li and H. Touyama. “Emotion Recognition Based on Group Phase Locking Value Using Convolutional Neural Network.” Scientific Reports 13 (2023): 3769. 10.1038/s41598-023-3769.

9) Dao, P. B. and W. J. Staszewski. “Lamb Wave Based Structural Damage Detection Using Stationarity Tests.” Materials (Basel) 14, no. 22 (November 12, 2021): 6823. 10.3390/ma14226823.

10) Daud, S. N. S. S. and R. Sudirman. “Wavelet-Based Filters for Artifact Elimination in Electroencephalography Signal: A Review.” Annals of Biomedical Engineering 50, no. 10 (October 2022): 1271–1291. 10.1007/s10439-022-03053-5.

11) Dressler, O., G. Schneider, G. Stockmanns and E. F. Kochs. “Awareness and the EEG Power Spectrum: Analysis of Frequencies.” BJA: British Journal of Anaesthesia 93, no. 6 (December 2004): 806–809. 10.1093/bja/aeh270.

12) Friedman, Nir, Tomer Fekete, Kobi Gal and Oren Shriki. “EEG-Based Prediction of Cognitive Load in Intelligence Tests.” Frontiers in Human Neuroscience 13 (June 11, 2019). 10.3389/fnhum.2019.00191.

13) Gage, N. M., J. Juranek, P. A. Filipek, K. Osann, P. Flodman, A. L. Isenberg and M. A. Spence. “Rightward Hemispheric Asymmetries in Auditory Language Cortex in Children with Autistic Disorder: An MRI Investigation.” Journal of Neurodevelopmental Disorders 1, no. 3 (September 2009): 205–214. 10.1007/s11689-009-9010-2.

14) García-Raffi, L. M. and E. Torrano. 2021. “Mittag-Leffler Functions and Their Applications in Fractional Calculus.” arXiv, March 23, 2021. https://arxiv.org/abs/2103.12559.

15) Guo, S., K. M. Kendrick, J. Zhang, M. Broome, R. Yu, Z. Liu and J. Feng. “Brain-Wide Functional Inter-Hemispheric Disconnection Is a Potential Biomarker for Schizophrenia and Distinguishes It from Depression.” NeuroImage: Clinical 2 (June 23, 2013): 818–826. 10.1016/j.nicl.2013.06.008.

16) He, K., Q. Hua, Q. Li, Y. Zhang, X. Yao, Y. Yang, W. Xu, J. Sun, L. Wang, A. Wang, G. J. Ji and K. Wang. “Abnormal Interhemispheric Functional Cooperation in Schizophrenia Follows the Neurotransmitter Profiles.” Journal of Psychiatry & Neuroscience 48, no. 6 (November–December 2023): E452–E460. 10.1503/jpn.230037.

17) Jaušovec, N. and K. Jaušovec. “Differences in Event-Related and Induced Brain Oscillations in the Theta and Alpha Frequency Bands Related to Human Intelligence.” Neuroscience Letters 293, no. 3 (2000): 191–94. 10.1016/s0304-3940(00)01526-3.

18) Jaušovec, N. and K. Jaušovec. “Differences in EEG Current Density Related to Intelligence.” Cognitive Brain Research 12, no. 1 (2001): 55–60. 10.1016/S0926-6410(01)00029-5.

19) Jaušovec, N. and K. Jaušovec. “Spatiotemporal Brain Activity Related to Intelligence: A Low Resolution Brain Electromagnetic Tomography Study.” Brain Research Cognitive Brain Research 16, no. 2 (2003): 267–72. 10.1016/s0926-6410(02)00282-3.

20) Jaušovec, N. and K. Jaušovec. “Sex Differences in Brain Activity Related to General and Emotional Intelligence.” Brain and Cognition 59, no. 3 (2005): 277–86. 10.1016/j.bandc.2005.08.001.

21) Jaušovec, N. and K. Jaušovec. “Emotional Intelligence and Gender: A Neurophysiological Perspective.” In Handbook of Individual Differences in Cognition, edited by A. Gruszka, G. Matthews and B. Szymura, 109–26. New York: Springer, 2010.

22) Miskovic, V. and A. Keil. “Reliability of Event-Related EEG Functional Connectivity during Visual Entrainment: Magnitude Squared Coherence and Phase Synchrony Estimates.” Psychophysiology 52, no. 1 (January 2015): 81–89. 10.1111/psyp.12287.

23) Monaim, H. and S. Fahlaoui. “General One-Dimensional Clifford Fourier Transform and Applications to Probability Theory.” Rendiconti del Circolo Matematico di Palermo, Serie II 73 (2024): 1453–66. 10.1007/s12215-023-00994-1.

24) Perkins, T. J., M. A. Stokes, J. A. McGillivray, A. J. Mussap, I. A. Cox, J. J. Maller and R. G. Bittar. “Increased Left Hemisphere Impairment in High-Functioning Autism: A Tract-Based Spatial Statistics Study.” Psychiatry Research: Neuroimaging 224, no. 2 (November 30, 2014): 119–123. 10.1016/j.pscychresns.2014.08.003.

25) Redwan, S. M., M. P. Uddin, A. Ulhaq and others. “Power Spectral Density-Based Resting-State EEG Classification of First-Episode Psychosis.” Scientific Reports 14 (2024): 15154. 10.1038/s41598-024-66110-0.

26) San-Segundo, R., Gil-Martín, M., D’Haro-Enríquez, L. F., and Pardo, J. M. 2019. “Classification of Epileptic EEG Recordings Using Signal Transforms and Convolutional Neural Networks.” Computers in Biology and Medicine 109 (June): 148–58. 10.1016/j.compbiomed.2019.04.031.

27) Saugstad, L. F. “A Lack of Cerebral Lateralization in Schizophrenia Is within the Normal Variation in Brain Maturation but Indicates Late, Slow Maturation.” Schizophrenia Research 39, no. 3 (October 19, 1999): 183– 196. 10.1016/s0920-9964(99)00073-0.

28) Sitnikova, E., Hramov, A. E., Koronovsky, A. A., and van Luijtelaar, G. 2009. “Sleep Spindles and Spike-Wave Discharges in EEG: Their Generic Features, Similarities and Distinctions Disclosed with Fourier Transform and Continuous Wavelet Analysis.” Journal of Neuroscience Methods 180 (2): 304–16. 10.1016/j.jneumeth.2009.04.006.

29) Tarasov, Vasily E. 2018. “Fractional Dynamics of Systems with Long-Range Interaction and Memory.” Frontiers in Physics 6: 110. 10.3389/fphy.2018.00110.

30) Thatcher, R.W., D. North and C. Biver. “EEG and Intelligence: Relations between EEG Coherence, EEG Phase Delay and Power.” Clinical Neurophysiology 116, no. 9 (September 2005): 2129–41.

31) Tozzi, A., E. Bormashenko and N. Jausovec. “Topology of EEG Wave Fronts.” Cognitive Neurodynamics 15 (2021a): 887–96. 10.1007/s11571-021-09668-z.

32) Tozzi, A., J. F. Peters, N. Jausovec, A. P. H. Don, S. Ramanna, I. Legchenkova and E. Bormashenko. “Nervous Activity of the Brain in Five Dimensions.” Biophysica 1, no. 1 (2021b): 38–47. 10.3390/biophysica1010004.

33) Turalska, Malgorzata and Bruce J. West. 2018. “Fractional Dynamics of Individuals in Complex Networks.” Frontiers in Physics 6: 110. 10.3389/fphy.2018.00110.

34) Yuan Q, Zhou W, Xu F, Leng Y, Wei D. Int J Neural Syst. 2018 Oct;28(8):1850010. doi: 10.1142/S0129065718500107. Epub 2018 Mar 19. PMID: 29665725

35) Zhang, Yan, Yuanhua Qiao, Lijuan Duan and Jun Miao. “Multistability of Almost Periodic Solution for Clifford-Valued Cohen–Grossberg Neural Networks with Mixed Time Delays.” Chaos, Solitons & Fractals 176 (November 2023): 114100. 10.1016/j.chaos.2023.114100.

